# Interleukin-4 restores neurogenic plasticity of the primary human neural stem cells through suppression of Kynurenic acid production upon Amyloid-beta42 toxicity

**DOI:** 10.1101/227306

**Authors:** Christos Papadimitriou, Hilal Celikkaya, Mehmet Ilyas Cosacak, Violeta Mashkaryan, Prabesh Bhattarai, Weilin Lin, Alvin Kuriakose Thomas, Yixin Zhang, Uwe Freudenberg, Carsten Werner, Caghan Kizil

**Affiliations:** German Center for Neurodegenerative Diseases (DZNE) Dresden, Helmholtz Association, Arnoldstr. 18, 01307, Dresden, Germany.; Center for Regenerative Therapies (CRTD), TU Dresden, Fetscherstr. 105, 01307, Dresden, Germany.; B CUBE, Center for Molecular Bioengineering, TU Dresden, Arnoldstr. 18, 10307, Dresden, Germany.; Leibniz Institute of Polymer Research Dresden, Max Bergmann Center of Biomaterials Dresden, Hohe Str. 6, 01069, Dresden, Germany.

**Keywords:** Human neural stem/progenitor cell, plasticity, Amyloid-beta42, Interleukin-4, KAT2, Kynurenic acid, Alzheimer’s disease

## Abstract

The immune response is an important determinant of the plasticity and neurogenic capacity of neural stem cells (NSCs) upon amyloid-beta42 (Aβ42) toxicity in Alzheimer’s disease (AD). However, the direct effects of individual immuno-modulatory effectors on NSC plasticity remain to be elucidated and are the motivation for reductionist tissue-mimetic culture experiments. Using starPEG-Heparin hydrogel system that provides a defined 3D cell-instructive neuro-microenvironment culture system, sustains high levels of proliferative and neurogenic activity of human NSCs, and recapitulates the fundamental pathological consequences of Amyloid toxicity upon Aβ42 administration, we found that the anti-inflammatory cytokine interleukin-4 (IL4) restores the plasticity and neurogenic capacity of NSCs by suppressing the Aβ42-induced kynurenic acid-producing enzyme kynurenine aminotransferase 2 (KAT2), which we also found to be upregulated in the brains of the AD model, APP/PS1dE9 mouse. Our transcriptome analyses showed that IL4 treatment restores the expression levels of NSC and cortical subtype markers. Thus, our dissective neuro-microenvironment culture revealed IL4-mediated neuroinflammatory crosstalk for human NSC plasticity and predicted a new mechanistic target for therapeutic intervention in AD.

## Introduction

The neurogenic capacity of the brain relies on the endogenous reservoir or transplanted population of neural stem cells (NSCs) that could be harnessed for neuronal repair during neurodegenerative diseases (Gage and Temple, 2013; Wyss-Coray, 2016). Therefore, it is fundamentally important to understand how NSCs can be made to contribute to neuronal regeneration and how they are affected by disease conditions.

Amyloid-beta42 (Aβ42) deposition and neurofibrillary tangles constitute the hallmark pathologies in Alzheimer’s disease (AD) (Bertram et al., 2010; Haass and Selkoe, 2007), which is the most prevalent neurodegenerative disease. In addition to neuronal survival and synaptic transmission, Aβ42 impairs NSC proliferation (He et al., 2013; Taupin, 2009; Tincer et al., 2016). Therefore, the human brain undergoes neurodegeneration and at the same time cannot replenish the lost neurons upon Aβ42 due to hampered NSC proliferation and neurogenesis. These combinatorial effects exacerbate the manifestation of the disease (Heneka et al., 2015; Nalbantoglu et al., 1997; Selkoe, 2002). Although pathogenic effects of Aβ42 in neurons are well-studied (Selkoe, 2002), little is known about how Aβ42 impinges on NSC proliferation and neurogenic capacity and how we can circumvent this reduction.

Animal models of AD suggest a multifaceted immuno-modulatory regulation of NSCs (Glass et al., 2010; Heneka et al., 2015; Kizil et al., 2015b; Kyritsis et al., 2012; Wyss-Coray, 2006). In general, pro-inflammatory signaling is believed to impair stem cell plasticity, while anti-inflammatory cytokines restore the homeostatic neurogenic ability in NSCs (Kizil et al., 2015b; Kokaia et al., 2012; Kyritsis et al., 2014; Schwartz et al., 2013). These effects of anti-inflammatory cytokines are thought to be taking place through the regulation of macrophages and resolution of the pro-inflammatory signaling, which is detrimental to stem cell plasticity (Carpentier and Palmer, 2009; Heneka et al., 2015; Wyss-Coray, 2006). However, since NSCs also recruit immune-related pathways, they could be directly responsive to the immune-milieu (Carpentier and Palmer, 2009), yet the direct effects of anti-inflammatory cytokines on NSC plasticity in AD is not well-studied because of the presence of multiple cell types, and the pleiotropy of individual factors that hinder the analyses of direct crosstalk mechanisms (Schwartz et al., 2013; Tincer et al., 2016). For instance, Interleukin-4 (IL4) was shown to be an anti-inflammatory signal by ameliorating or exacerbating the amyloid-load and neuropathology in a context-dependent manner in mammals, and little is known about its direct effect on NSCs (Chakrabarty et al., 2012; Kiyota et al., 2010; Park et al., 2008). Recently, we have shown that IL4 establishes neuro-immune crosstalk between the site of pathology and NSCs, and directly regulates stem cell proliferation in an adult zebrafish AD model (Bhattarai et al., 2016). Therefore, investigating whether anti-inflammatory cytokine IL4 may have a similar mechanism of action in mammalian AD models and identifying the downstream regulation of immune system in NSCs would be instrumental in elaborating on the neuro-immune regulation of NSC plasticity. However, mouse models may not recapitulate the whole spectrum of the AD (LaFerla and Green, 2012) and the specific molecular programs in human cells differ from mouse cells (Qiu et al., 2016). Additionally, conventional 2D cultures are not representative of in vivo environments (Ravi et al., 2015) and emerging 3D culture technologies, such as organoids, are often highly complex. Therefore, alternative in vitro systems utilizing human cells to model neural development or neurodegeneration in a thoroughly defined and tunable in vivo-like 3D environment would be highly beneficial. The widely used Matrigel^TM^-based 3D cultures are of ill-defined composition, cannot be adjusted for various important cell-instructive parameters (Ravi et al., 2015) and are therefore not suitable for elucidating the influences of different exogenous and paracrine signals on cellular development (Choi et al., 2014; Fatehullah et al., 2016; Tang-Schomer et al., 2014; Zhang et al., 2014), which, in turn, limits their applicability for the investigation of the plasticity and neurogenic capacity of NSPCs. Therefore, reductionist, humanized assays of NSC plasticity and neurogenesis could be instrumental for elucidating the role of individual cytokines in vitro.

## Results

In this study, we utilized a modular platform of minimalist GAG-based hydrogels to generate defined tissue-mimetic neuro-microenvironments for testing the plasticity of NSCs and the network formation of neurons as well as the molecular aspects of neurodegenerative diseases. The choice of compositionally and biophysically defined GAG-containing materials that are known to enable the tissue-analogous presentation of molecular immunomodulators including IL4 (Lohmann et al., 2017; Schirmer et al., 2016) adds valuable new options to those of previously reported 3D culture models (Choi et al., 2014; Zhang et al., 2014).

The anti-inflammatory cytokine interleukin-4 (IL4) has previously been implicated in regulating NSC proliferation in rodent models indirectly by converting macrophages into a post-inflammatory state, implying a negative role for inflammation on stem cell plasticity (Griffin, 2013). Conversely, we have recently reported direct effects of IL4 on NSCs in an adult zebrafish brain model of AD, suggesting a positive role for pro-inflammatory cues in regeneration (Bhattarai et al., 2016). For humans, however, the direct role of IL4 on NSC plasticity and neurogenic capacity is still unknown. To address this question, we used the biohybrid starPEG-GAG hydrogel-based NSC cultures that are particularly well-suited to our question since the materials were previously shown to reversibly bind and protect IL4 in ways resembling its complexation in extracellular matrices (Schirmer et al., 2016), resulting in the effective modulation of the activity of anti-inflammatory cytokines (Freudenberg et al., 2016). To determine the effects of IL4 on our 3D NSC cultures in normal and AD conditions, we first analyzed the expression of interleukin-4 receptor (IL4R) and found that IL4R is expressed in GFAP-positive NSCs from the beginning of the culture and can lead to phosphorylation of STAT6 after IL4 treatment (Figure 1A-B’), confirming functional intracellular signaling. To determine whether IL4 treatment affects the normal development and composition of the human NSC cultures, we compared the control and IL4-treated cultures (Figure 1C-H). We observed that IL4 does not affect the total number and composition of NSCs and neurons in the gels (Figure 1I), indicating that IL4 does not alter the plasticity and neurogenic capacity of NSCs in neurodevelopmental process in 3D gels.

**Figure 1.**
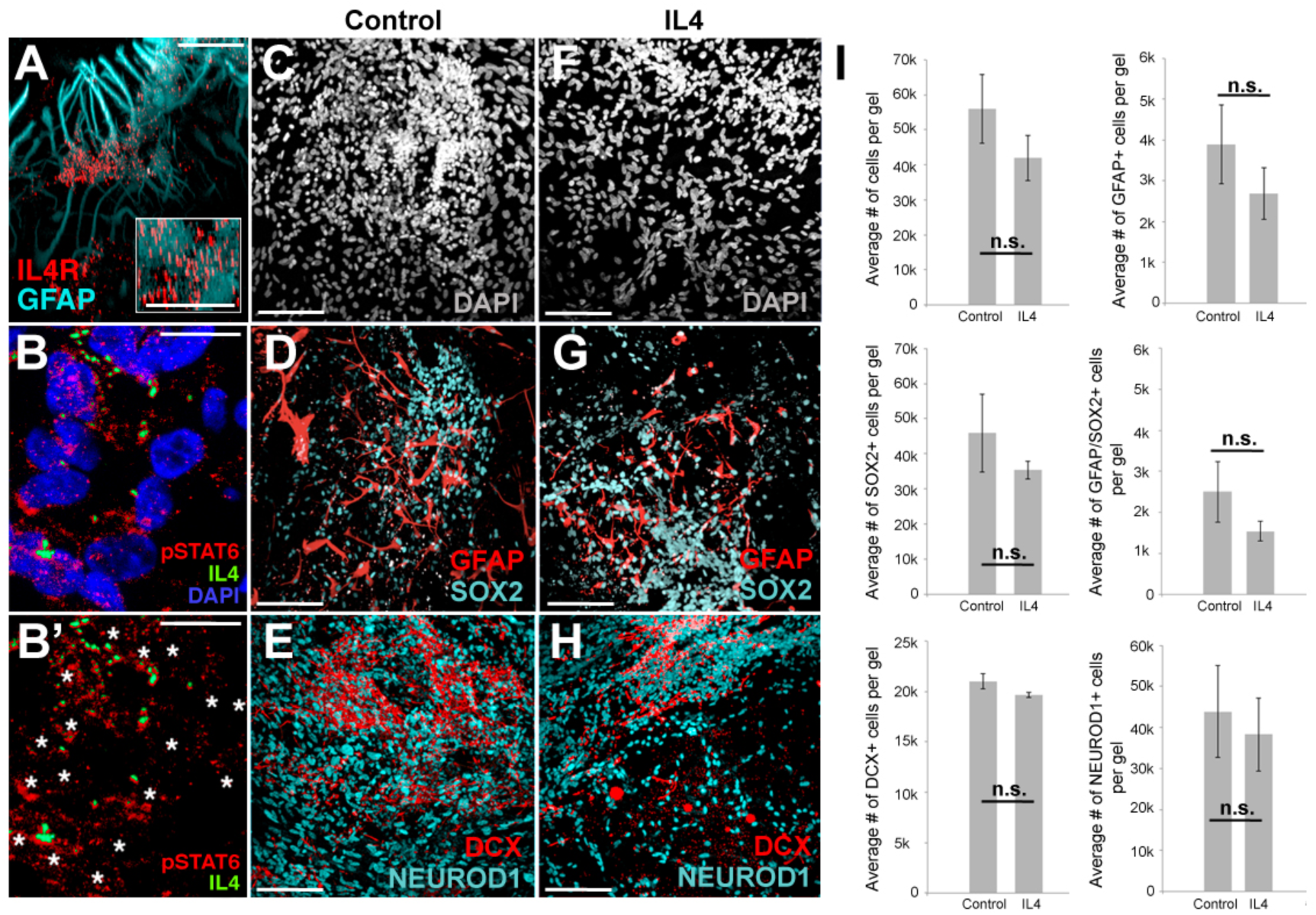
Effects of IL4 on 3D cultures of primary human NSCs. (A) IL4R in GFAP-positive glia. (B) pSTAT6 and IL4 in control gels. (B’) DAPI removed from B. Asterisks: nuclear pSTAT6. (C, F) Nuclei of the cells in control (C) and IL4-treated gels (F) shown in X-axis. (D, G) GFAP and SOX2 in control (D) and IL4-treated gels (G) shown in X-axis. (E, H) DCX and NEUROD1 in control (E) and IL4-treated gels (H) shown in X-axis. (I) Quantification of C-H. Scale bras: 25 *μ*m (inset in A, and B), and 100 *μ*m elsewhere. All gels are 3 weeks of culture. See also Supplementary Figure 1.

Previously, we have shown that in 3D gel cultures, primary human NSC plasticity, neurogenic ability, and neuronal network forming capacity are reduced by Aβ42 (Papadimitriou et al., 2017), and IL4 has a positive effect on NSC proliferation in an Aβ42-based Alzheimer’s model of adult zebrafish brain (Bhattarai et al., 2016). Therefore, to determine whether IL4 treatment would have an effect on the reduction of NSC plasticity, neurogenic properties and network formation of the neurons, we cultured Aβ42-treated cells in the presence of hydrogel-administered IL4, and compared it to control and Aβ42-treated cells without IL4 (Figure 2A-F). Compared to control gels, Aβ42 conditioning significantly reduced NSCs (GFAP/SOX2) and early neurons (NEUROD/DCX), while IL4 rescued this reduction (Figure 2G). Quantifying the activated (GFAP+/SOX2+) fraction of NSCs (GFAP and/or SOX2-positive cells), we found that IL4 significantly increased the percentage of activated NSCs (Figure 2H), which manifests itself in the formation of more DCX-positive neurons and networks (Figure 2I-I’’). To validate the positive effect of IL4 on activation of proliferation of NSCs and neurogenesis, we determined the levels of newborn cells at 3 weeks of 3D cultures after a 6-hour BrdU treatment during the first week (Figure 2J-L). We found that Aβ42 administration reduced the total number of newborn cells (˜96.1%) and BrdU+ glia significantly (˜86.5%), while IL4 treatment rescued these reductions and enhanced the ratio of newborn cells to the BrdU-positive GFAP cells (as an indicator of neurogenic capacity) (Figure 2M). These results indicate that Aβ42 impairs NSC plasticity, neurogenic capacity and network-formation ability of human NSCs and neurons while IL4 restores these features despite the prevalent AD environment (Figure 2N). We found that the rescue effect of IL4 is specific because knocking-down IL4 activity using a neutralizing antibody significantly reduced the rescue effect (Supplementary Figure 1). Overall, these results suggest that our starPEG-heparin 3D hydrogel cultures of human NSCs can recapitulate the tissue-mimetic manifestation of neurogenic capacity and plasticity and can be used to investigate the direct effects of particular immune-related factors in a highly reductionist manner.

**Figure 2.**
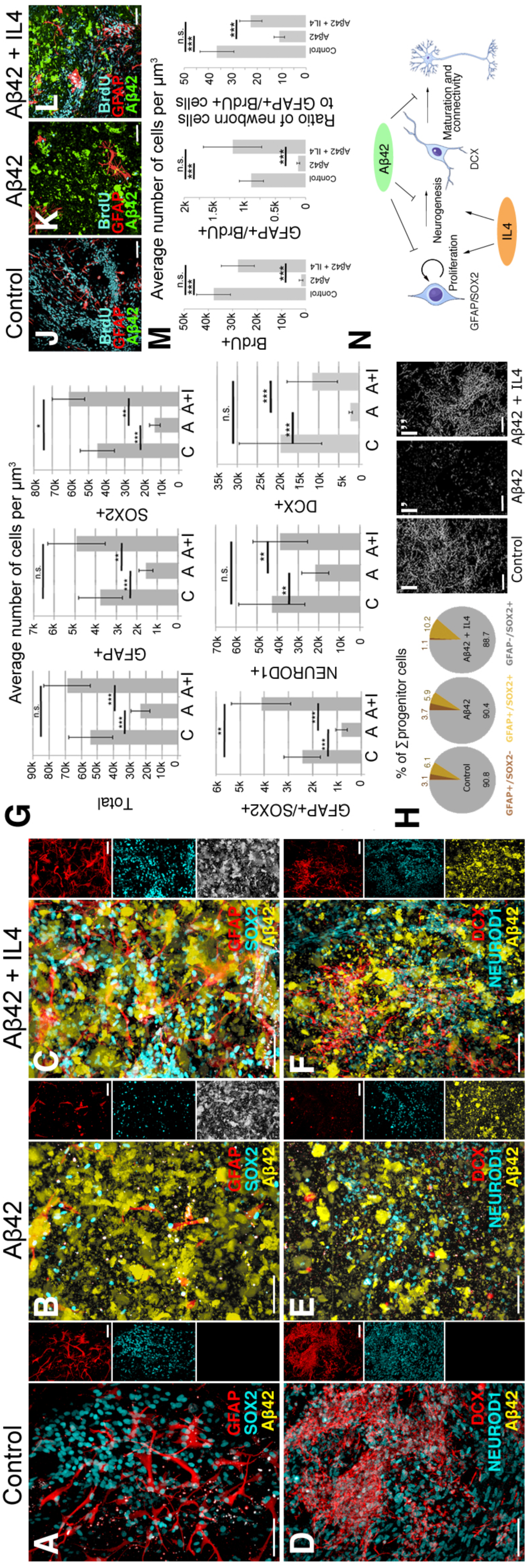
Effects of IL4 on the Aβ42 toxicity model in 3D cultures. (A-C), GFAP, SOX2, Aβ42 in control (A), Aβ42 (B) and Aβ42+IL4 gels (C). (D-F) DCX, NEUROD1, Aβ42 in control (D), Aβ42 (F) and Aβ42+IL4 gels (G). Small panels on the right side of A-F are single fluorescent channels. (G) Quantification graphs. (H) Pie-chart for composition of neural stem/progenitor cells. (I-I”) Skeletonized connected networks of DCX-positive neurons in (F-H). (J-L) BrdU, GFAP and Aβ42 in control (L), Aβ42 (M) and Aβ42+IL4 gels (N). (M) Quantification of (L-N). (N) Schematics for the effects of IL4 on the AD model. Scale bars: 50 *μ*m. All gels: 3 weeks of culture. See Supplementary Figure 2.

Since our 3D cultures can be used to investigate the direct effects of IL4 on NSCs, we also aimed to analyze the downstream regulation exerted by IL4. We previously found that IL4 increases stem cell proliferation in the adult zebrafish brain after Aβ42 administration (Bhattarai et al., 2016). Whole genome transcriptome analysis of this model revealed differentially expressed and enriched components of the tryptophan metabolism pathway that ultimately generated kynurenic acid (KYNA) (Bhattarai et al., 2016). While KYNA was reported to be a neuroprotective molecule (Schwarcz et al., 2012; Szalardy et al., 2012; Zwilling et al., 2011), its direct effect on NSCs is unknown (Jones et al., 2013). We hypothesized that the effects of IL4 on Aβ42-mediated impairment of NSC plasticity and neurogenic capacity would regulate KYNA production.

To test this hypothesis, we compared the expression of the enzymes producing KYNA, which is produced from tryptophan by a cascade of enzymatic reactions involving three main enzymes: IDO1, TDO2 and KAT2 (Figure 4A-D, Supplementary Figure 2). We found that the percentage of cells expressing IDO1 and TDO2 remained constant after Aβ42 or IL4 treatment (Supplementary Figure 2G). However, Aβ42 increased the levels of KAT2 (Figure 3E), which is expressed in glial cells as described previously (Schwarcz et al., 2012). IL4 restored the original percentage of cells expressing this enzyme (Figure 3E), suggesting that the toxic effect of Aβ42 on NSCs is, in part, mediated by the upregulation of KAT2 and the production of KYNA. Therefore, we hypothesized that an effective concentration of produced in cultures should correlate with Aβ42 toxicity, and IL4 treatment could reduce these levels. To test this hypothesis, we performed mass spectrometry coupled with liquid chromatography for detecting the levels of KYNA in cell culture medium from control, IL4-treated, Aβ42-treated, and Aβ42 + IL4-treated gels during their last week of culture (Supplementary Figure 3). We found that the amount of KYNA produced by GFAP cells was significantly increased after Aβ42 treatment, and IL4 reduces this level down to control levels (Figure 3E’, Supplementary Figure 3D). These results indicate that KYNA mediates Aβ42 toxicity in human NSCs, and IL4 reduces effective KYNA levels to physiological levels.

**Figure 3.**
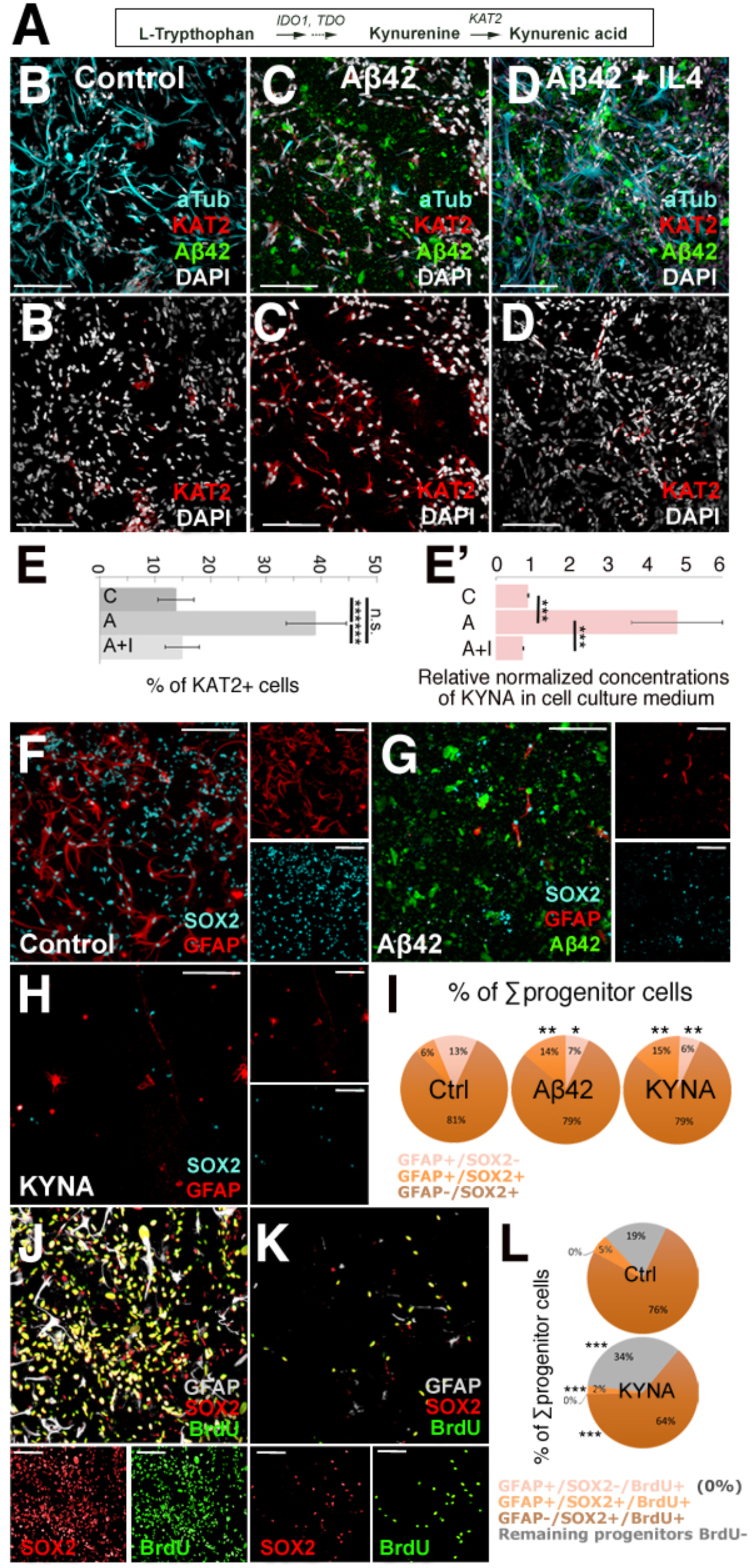
Aβ42 toxicity is mediated by Kynurenic acid production by KAT2. (A) Tryptophan metabolism of kynurenic acid. (B-D’) Acetylated-tubulin, KAT2, Aβ42 in control (B), Aβ42-treated (C), and Aβ42+IL-treated gels (D). KAT2 is shown alone in (B’-D’). (E) Quantification of the percentage of cells expressing KAT2. C: control, A: Aβ42, A+i: Aβ42+IL4. (E’) Quantification of KAT2 levels. C: control, A: Aβ42, A+i: Aβ42+IL4. (F-H) SOX2 and GFAP in control (F), Aβ42-treated (G), 10 μM KYNA-treated (H) gels. Single fluorescent channels are on the right. (I) Composition of progenitors as percentage. (J,K) GFAP, SOX2, and BrdU in control (J) and KYNA-treated (K) gels. (L) Composition of proliferating progenitors as percentage. Scale bars: 100 *μ*m. All gels: 3 weeks of culture. See also Supplementary Figure 3 and 4.

To investigate how KYNA affects NSCs in 3D cultures and whether this effect is similar to that of Aβ42 treatment, we performed immunocytochemical staining for GFAP and SOX2 on control, Aβ42-treated, and KYNA-treated NSPC cultures (Figure 3F-H). We found that KYNA reduced the total number of GFAP+, SOX2+ and GFAP/SOX2 double-positive cells similar to Aβ42 (Supplementary Figure 4A) and reduced the percentage of activated NSCs (GFAP+/SOX2+) (Figure 3I, Supplementary Figure 4A). Furthermore, compared to control cultures, KYNA diminished the proliferative capacity of NSCs (Figure 3J-L, Supplementary Figure 4B-D). This result shows that KYNA is an intermediate conveying the Aβ42-induced impairment of the proliferative capacity of NSCs.

**Figure 4.**
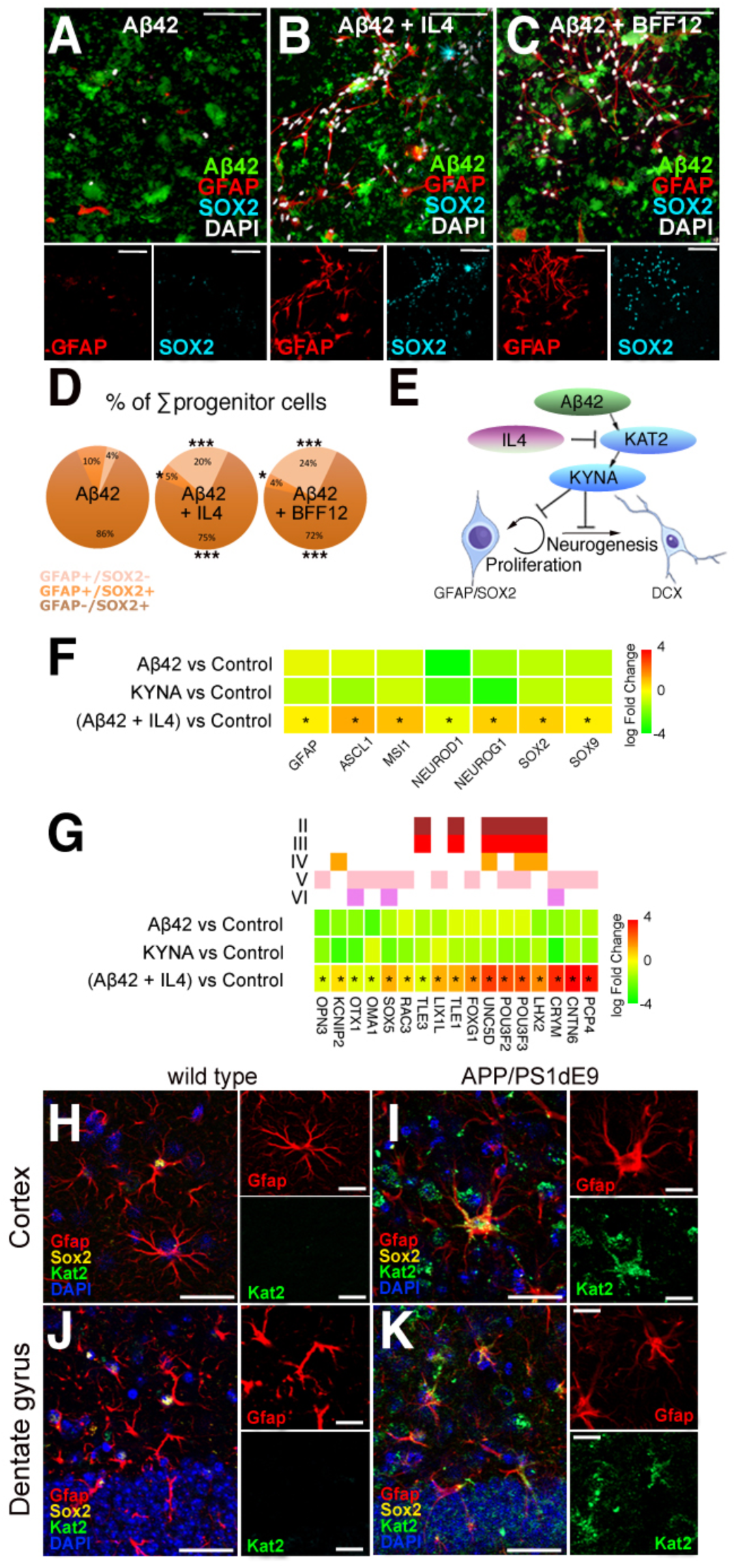
IL4 rescues KYNA-mediated Aβ42 toxicity on primary human NSCs. (A-C) GFAP and SOX2 in Aβ42-treated (A), Aβ42+IL4-treated (B), Aβ42+BFF12-treated gels (C). Lower panels are individual channels. (D) Composition of progenitor cells as percentage. (E) Schematics for the functional interaction of Aβ42, KAT2, KYNA and IL4 in regulating human NSPC plasticity and neurogenic capacity during AD. (F) Heat map for expression changes of genes related to NSC plasticity in Aβ42, KYNA, and Aβ42 + IL4 conditions. (G) Heat map for expression changes of cortical subtype marker genes in Aβ42, KYNA, and Aβ42 + IL4 conditions. (H-K) Gfap, Sox2, Kat2 in the 12-month-old control (H, J) and APP/PS1dE9 (I, K) AD model mouse in the cortex and dentate gyrus. Scale bars: 100 *μ*m. All gels: 3 weeks of culture. See also Supplementary Figure 4.

Since IL4 reduces the production of KYNA by suppressing the expression of KAT2, we hypothesized that blocking the KAT2 activity would mimic the effects of IL4 on NSC plasticity. Therefore, we inhibited KAT2 with the selective antagonist BFF12, and found that the reduction in GFAP+ and SOX2+ cells by Aβ42 is counteracted by BFF12 treatment similar to IL4 (Figure 4A-C). Thus, the restorative effect of KAT2 inhibition by BFF12 on the diminished NSCs (GFAP/SOX2) is comparable to IL4-treatment (Figure 4D), confirming that kynurenic acid production is one reason for Aβ42 toxicity in NSCs. Overall, we showed that Aβ42 reduces human NSC proliferation and neurogenic capacity in dissective AD model conditions in GAG-based hydrogels through upregulation of KAT2 and subsequent increases in kynurenic acid, which can be prevented by IL4 through the inhibition of KAT2 (Figure 4E).

Based on our findings, we hypothesized that if IL4 can restore the plasticity and neurogenic output of human NSCs in Aβ42 toxicity conditions mediated by KYNA, the expression of NSC makers and cortical markers should change similarly in Aβ42 and KYNA-treated gels, and IL4 treatment should restore those expression levels. Therefore, we performed whole transcriptome sequencing on gels treated with Aβ42, KYNA, and Aβ42 with IL4. We found that in Aβ42- or KYNA-treated gels, there is an overall reduction in NSC marker expression (Figure 4F, upper and middle rows), while IL4 treatment with Aβ42 abrogates this reduction and in some cases even enhances the expression levels of NSC markers (Figure 4F, lower row). Since NSC marker expression levels are restored by IL4 treatment after Aβ42, we hypothesized that this change in NSCs might be reflected in the replenishment of cortical subtypes. Therefore, we analyzed a set of cortical subtype markers (Figure 4G) and observed that similar to NSC markers, Aβ42 and KYNA treatments reduce the cortical marker expression in general (upper and middle row, Figure 4G), while IL4 restores or enhances the expression levels of cortical neuronal markers (Figure 4G, lower row). These results suggest that IL4 restores the neurogenic ability of NSCs and neuronal network formation after Aβ42 toxicity through restoring the molecular programs that underlie the NSC plasticity and neurogenic output.

KAT2 is expressed in a subset of astrocytes in the cerebral cortex and hippocampus of rat brains (Guidetti et al., 2007); however, its regulation by Aβ42 conditions and pathology is unknown. Therefore, to test whether the findings in our GAG-based hydrogel system would be biologically relevant to the in vivo situation, we analyzed the expression of KAT2 in controls and APP/PS1dE9 model of AD mouse brains (Figure 4F-I). Compared to cortical and hippocampal regions of control animals where KAT2 is detected rather weakly in very few cells (Figure 4H, J), AD mouse brains strongly upregulated KAT2 levels in GFAP-positive glia in the cortex and the hippocampus (Figure 4I, K). These results support our findings that Aβ42 toxicity in NSCs is mediated by kynurenic acid through upregulation of KAT2, and indicates that our 3D culture system can be used as a predictive tool for in vivo conditions.

## Discussion

KYNA was previously shown to have a neuroprotective role in neurodegenerative diseases (Klein et al., 2013; Schwarcz et al., 2012; Stone and Darlington, 2002; Szalardy et al., 2012; Zwilling et al., 2011), but its role in NSCs was not clear. Here, we demonstrated that KYNA negatively affects human NSC plasticity in a dissective, tissue-mimetic 3D culture system. This finding is important because clinical efforts for enhancing KYNA levels might be effective for neuronal survival but could impair NSC activity that is required to replenish the lost neurons. Our data imply that a temporal control of KYNA production could be beneficial by initially enhancing NSC proliferation and later sustaining neuroprotection. Furthermore, we showed that IL4 has, in addition to its documented neuromodulatory role, a direct effect on human NSCs by a previously unknown involvement in KAT2 expression and KYNA production. Additionally, these results suggest that our 3D culture system can be used to pinpoint previously unidentified roles of highly studied molecules during neurodegenerative diseases. For instance, IL4 is an anti-inflammatory factor, and its expression is contingent upon advanced AD pathology, when amyloid load and plaques already exist (Heneka et al., 2015; Schwartz et al., 2013). Previously, the beneficial effects of IL4 on neuronal survival and NSC activity were associated with the reduced inflammatory milieu (Kiyota et al., 2010). However, our dissective GAG-based hydrogel 3D cultures are devoid of an immune system. Therefore, our data indicated that IL4 establishes direct crosstalk between the immune system and the NSC compartment, governing plasticity and the neurogenic capacity of stem cells. This finding not only confirms the previous in vivo results in zebrafish (Bhattarai et al., 2016) but also provides validation for the use of GAG-based hydrogel culture as an experimentally reliable and reductionist surrogate for in vivo studies. Our 3D cultures can also open up new avenues for tweaking the neuroinflammatory microenvironment toward therapeutically relevant mobilization schemes of endogenous NSCs.

Our newly established methodology of GAG-based hydrogel NSC cultures was key to the reported analyses of the effects of IL4 and KYNA since this reductionist 3D neuro-microenvironment system concomitantly supported NSC plasticity and neurogenic potential. As shown for the specific reported case, our tissue-mimetic reductionist culture system is highly advantageous for elucidating the effects of individual immunomodulators on NSCs, for recapitulating complex micromilieu, and for biologically investigating the downstream effects of a particular signaling pathway. In particular, the GAG-based hydrogel-culture approach is similarly suitable for investigating the cellular interactions of neuroglia with macrophages, i.e., elucidating the interplay between cellular components of the immune system and NSCs in AD mimicking conditions. The GAG-based hydrogel culture system also has the potential to facilitate high-throughput screening of biologically active compounds for their effects on NSC plasticity and specific neuro-immune communication.

## Author contributions

C.P. and C.K. conceived and designed the experiments. L.B., U.F., and C.W. provided the gel materials, C.P. performed cell cultures, imaging and quantifications. P.B., M.I.C., H.H., H.C. and V.M helped the cell cultures. Y.Z. provided the Amyloid peptide, W.L. performed LC-MS/MS. C.K. wrote the manuscript, C.K. and C.W. revised the manuscript.

## Acknowledgements

This work was supported by DZNE and Helmholtz Association (VH-NG-1021, C.K.), DFG (KI1524/6, C.K.), (AN797/4-1, C.L.A.) and (CRC TR 67, CRC SFB 655, FOR/EXC999, C.W.); and BMBF (PRECIMATRIX-FKZ-03XP0083–310117, C.W.).

## Materials and Methods

### Generation of starPEG-Heparin hydrogels and synthesis of Amyloid peptides

StarPEG-heparin hydrogels were generated as previously described (Maitz et al., 2013; Wieduwild et al., 2013) and (Papadimitriou et al., 2017). Amyloid peptides were synthesized as previously described (Bhattarai et al., 2017; Bhattarai et al., 2016; Kizil et al., 2015a; Wieduwild et al., 2013).

### Primary human neural stem cell cultures and treatments

Primary neural stem cells isolated from the cerebral cortex at gestation week 21 were obtained from ScienCell Research Laboratory (SRL, Catalog Number 1800, Carlsbad, CA, USA) at passage one and delivered as frozen stocks. The cells are certified to be negative for HIV-1, HBV, HCV, mycoplasma, bacteria, yeast, and fungi. NSCs were seeded on conventional T75 flasks or 24-well plates and cultured with Astrocyte medium (SRL, Catalog Number 1801) supplemented with 2% fetal bovine serum (SRL, Catalog Number 0010), 1% astrocyte growth supplement (SRL, Catalog Number 1852) and 1% penicillin/streptomycin solution (SRL, Catalog Number 0503) in an incubator with a 5% CO2/95% air atmosphere at 37 °C. BrdU was added to the culture medium at 1 week after encapsulation (10 mg/ml) for 1 day. KYNA, IL4 and BFF12 were present in the culture medium throughout the culture at the following concentrations: IL4 (10 nM), KYNA (10 uM), BFF12 (10 uM).

### Immunocytochemistry

All of the hydrogels were fixed with ice-cold 4% paraformaldehyde and incubated for 1.5 hours at room temperature followed by washing in PBS overnight at 4 °C. For immunocytochemistry, the hydrogels were blocked and permeabilized in blocking solution for 4 hours at room temperature. For BrdU-treatment, the gels were incubated with 2 M HCl for 20 minutes at 37 °C followed by three washes in PBS (2 hours each). EdU staining was performed according to the manufacturer’s protocol (Life Technologies, C10638) using a 1 hour incubation step. The hydrogels were incubated with primary antibodies (Supplementary Table 1) in blocking solution overnight at 4 °C. The gels were washed for two subsequent days at 4 °C, with occasional changes of the PBS. After washing, the gels were incubated with the secondary antibodies (1:500 in blocking solution) at room temperature for 6 hours. After 3 washing steps of 2 hours each, DAPI staining was performed (1:3000 in PBS, 2 hours at room temperature). Immunostaining for SOX2 (Santa Cruz Biotechnology, 1:100), TUBB3 (R&D Systems, 1:500), GFAP (Novex, 1:500), DCX (Novex, 1:300), Aβ42 (Cell Signaling Technology, host: Rabbit, 1:500), BrdU (AdB Serotec, 1:500), TDO2 (Novus Biologicals, 1:300), IDO1 (Novus Biologicals, 1:300), KAT2 (Sigma, 1:300) was performed. All of the secondary antibodies were conjugated to AlexaFluor dyes (Life Technologies).

### Fluorescent imaging

For the hydrogels, fluorescent imaging was performed using a Leica SP5 inverted Laser Scanning Confocal microscope. The hydrogels were placed in glass bottom Petri dishes. Sixty microliters of PBS were added on top of the hydrogels to avoid desiccation. The Z-stacks were captured using a 25x water immersion lens. Every Z-stack had a z-distance of 500 μm. Monolayers were imaged using an inverted Zeiss Apotome 2 microscope.

### Tandem mass spectroscopy coupled to liquid chromatography (LC-MS/MS)

For LC-MS/MS, the culture media in last week (Day 15-21) were collected during the medium change for untreated (control), Aβ42-treated, IL4-treated, and Aβ42+IL4-treated gels. In total, 10 ml of medium for every condition from 6 gels per experimental group was collected. The quantification of Kynurenic acid (KYNA) was performed by tandem mass spectrometry (MS/MS) coupled to liquid chromatography (LC-MS/MS) using waters ACQUITY UPLC system with ACQUITY TQ Detector. The UPLC was equipped with an analytical C18 column (ACQUITY UPLC ^®^ BEH C18 1.7 *μ*m, 2.1 x 50 mm). The samples with 10 *μ*L of the volume were injected to the column. To avoid the contamination between the samples, 3 injections of PBS was performed after each sample measurement. Water with 0.1% formic acid was used as solvent A and acetonitrile with 0.1% formic acid as solvent B. The time curve for running: 0.0-1.0 min, 100 % buffer A; 1.0-4.0 min, a linear gradient running from 100 % buffer A to 100 % buffer B; 4.0 min to 5.0 min, 100 % buffer B; 5.0 min to 5.5 min, a linear gradient running from 100 % buffer B to 100 % buffer A; 5.5 min to 6.0 min, 100 % buffer A. For MS/MS analysis of KYNA, the TQ detector was set in multiple reaction monitoring (MRM) to detect the parent (189.95 m/z) and the daughter (88.98 m/z), with 2770V of the capillary voltage, 32V of the cone energy, 40V of the collision energy and 0.328s of the dwell time.

### Next-generation sequencing and bioinformatics analyses

Sample preparation, RNA isolation, library preparation, next-generation sequencing, data analyses are performed as described before (Papadimitriou et al., 2017).

### APP/PS1dE9 mouse model of Alzheimer’s disease

APP/PS1dE9 mouse brains were gifts from Gerd Kempermann. Immunohistochemistry on paraffin-embedded sections were performed as described before (van Praag et al., 1999).

### Image analysis and statistics

The 3D reconstructions of hydrogel images and videos were generated using Arivis 4D software. Images from monolayers were processed using Zeiss ZEN software. The statistical analyses were performed using GraphPad Prism and two-tailed Student’s t-tests. The levels of significance were *: p ≤ 0.05, **: p ≤ 0.01, and ***: p ≤ 0.001. In all graphs, means ± standard deviations are shown. Skeletonized networks are constructed as described (Papadimitriou et al., 2017).

The effect size was calculated using G-Power, and the sample size was estimated with n-Query. The data conforms to normal distribution as determined by Pearson’s chi-squared test. The variations between the samples are similar as determined by variance estimation using Microsoft Excel software. For 3D gels, 9 gels were used for quantifications (3 technical replicates in every experiment, and 3 experiments as biological replicates). All experiments were replicated many times in the laboratory and results were confirmed independently (80-120 gels were qualitatively analyzed to check the consistency of the results for every individual experiment).

## Supplementary Information

**Supplementary Figure 1.**
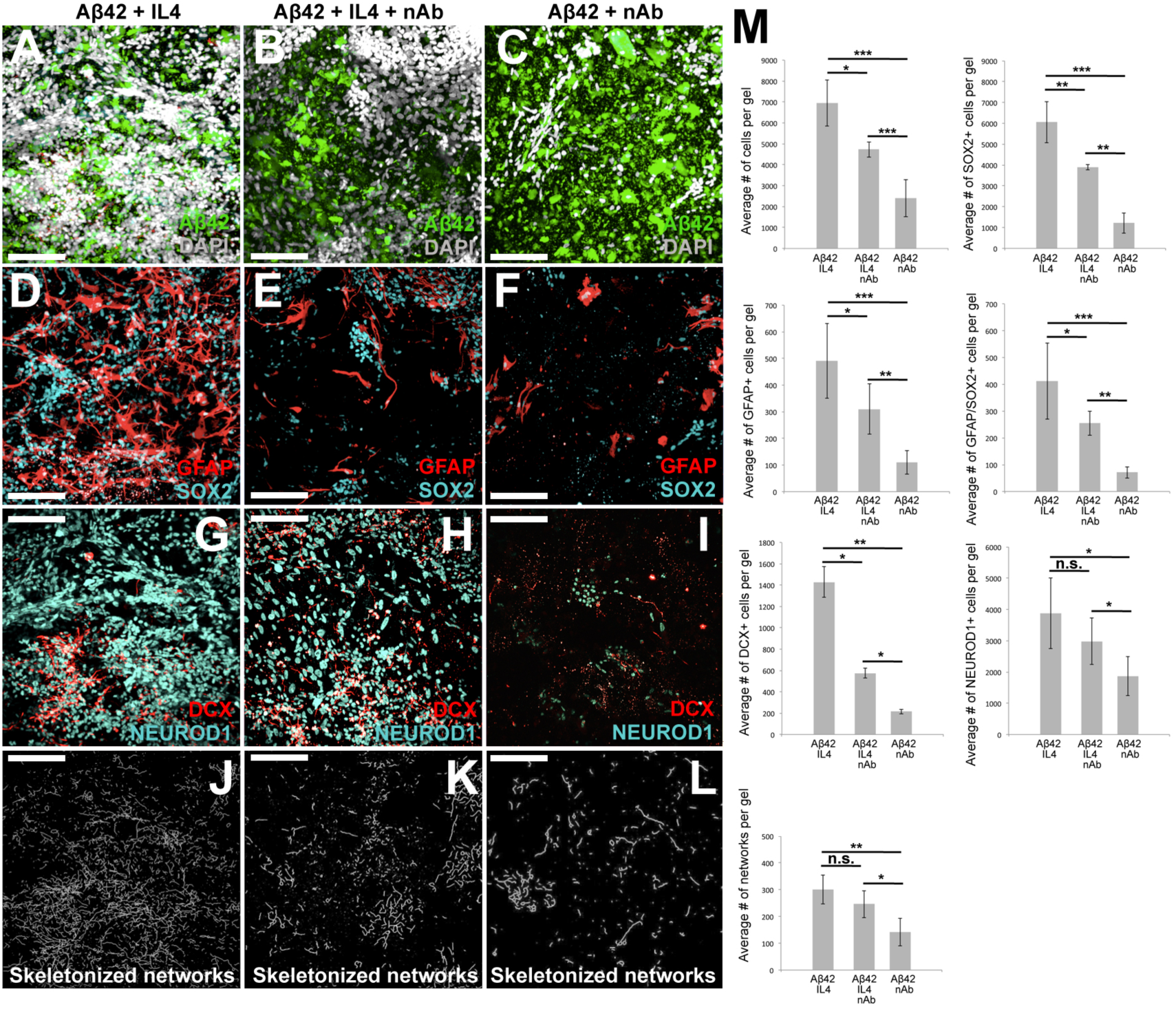
The effects of IL4 on primary human NSCs are specific. (A-C) Aβ42 and DAPI in Aβ42+IL4-treated (A), Aβ42 + IL4 + neutralizing IL4 antibody (nAb)-treated (B), and Aβ42 + nAb-treated gels (C). (D-F) GFAP and SOX2 in Aβ42+IL4-treated (D), Aβ42 + IL4 + nAb-treated (E), and Aβ42 + nAb-treated gels (F). (G-I) DCX and NEUROD1 in Aβ42+IL4-treated (G), Aβ42 + IL4 + nAb-treated (H), and Aβ42 + nAb-treated gels (I). (J-L) Skeletonized networks of connected paths in Aβ42+IL4-treated (J), Aβ42 + IL4 + nAb-treated (K), and Aβ42 + nAb-treated gels (L). (M) Quantifications of A-L. Scale bars: 100 *μ*m. All gels are 3 weeks of culture. Related to Figure 1.

**Supplementary Figure 2.**
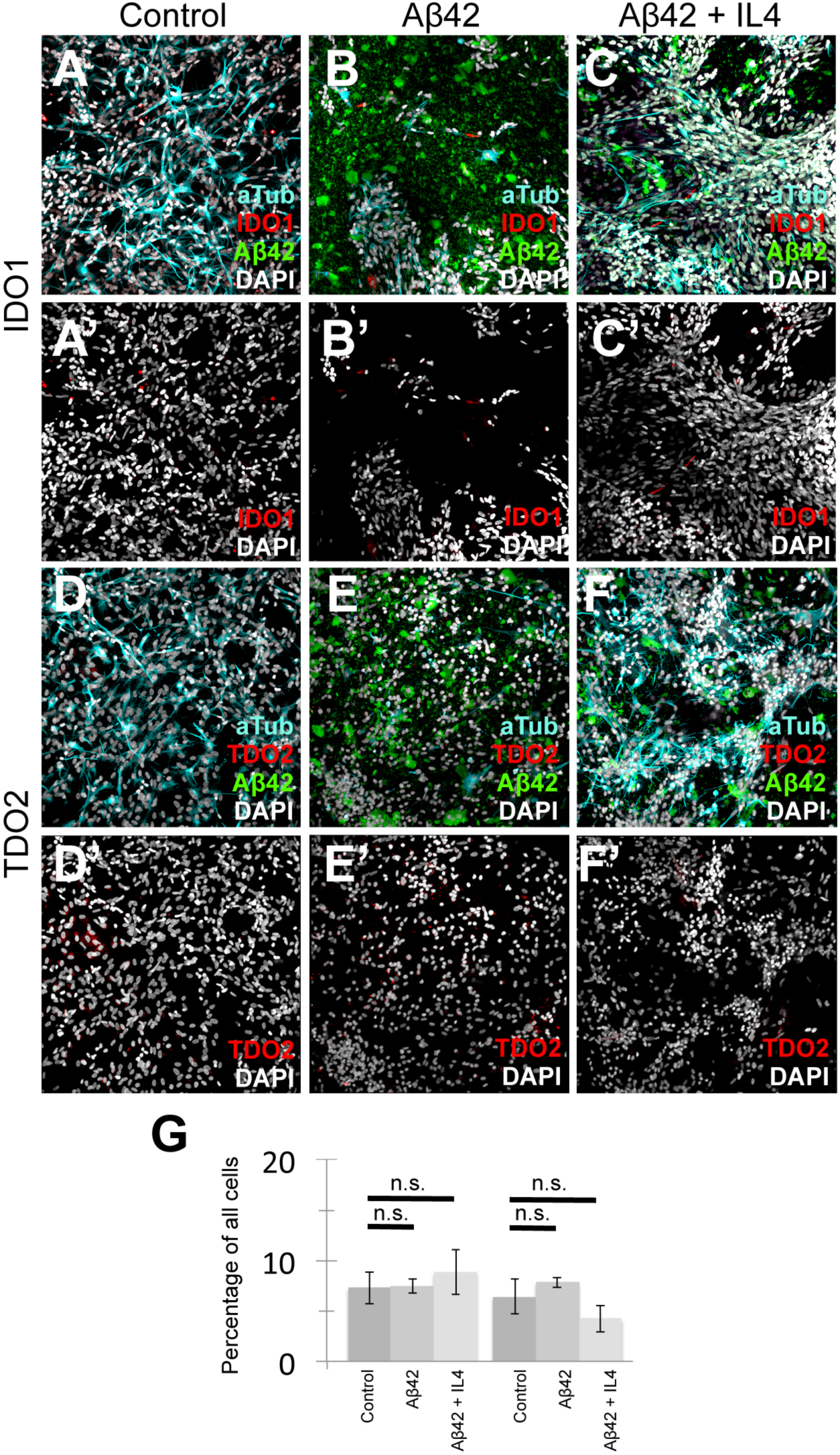
Expression of TDO and IDO1. (A-C) Acetylated tubulin, IDO1 and Aβ42 in control (A), Aβ42-treated (B), Aβ42 + IL4-treated gels (C). (A’-C’) IDO1 and DAPI channels of A-C, respectively. (D-F) Acetylated tubulin, TDO and Aβ42 in control (D), Aβ42-treated (E), Aβ42 + IL4-treated gels (F). (D’-F’) TDO and DAPI channels of D-F, respectively. (G) Quantification graph for A-F’. Scale bars: 100 *μ*m. All gels are 3 weeks of culture. Related to Figure 2 and 3.

**Supplementary Figure 3.**
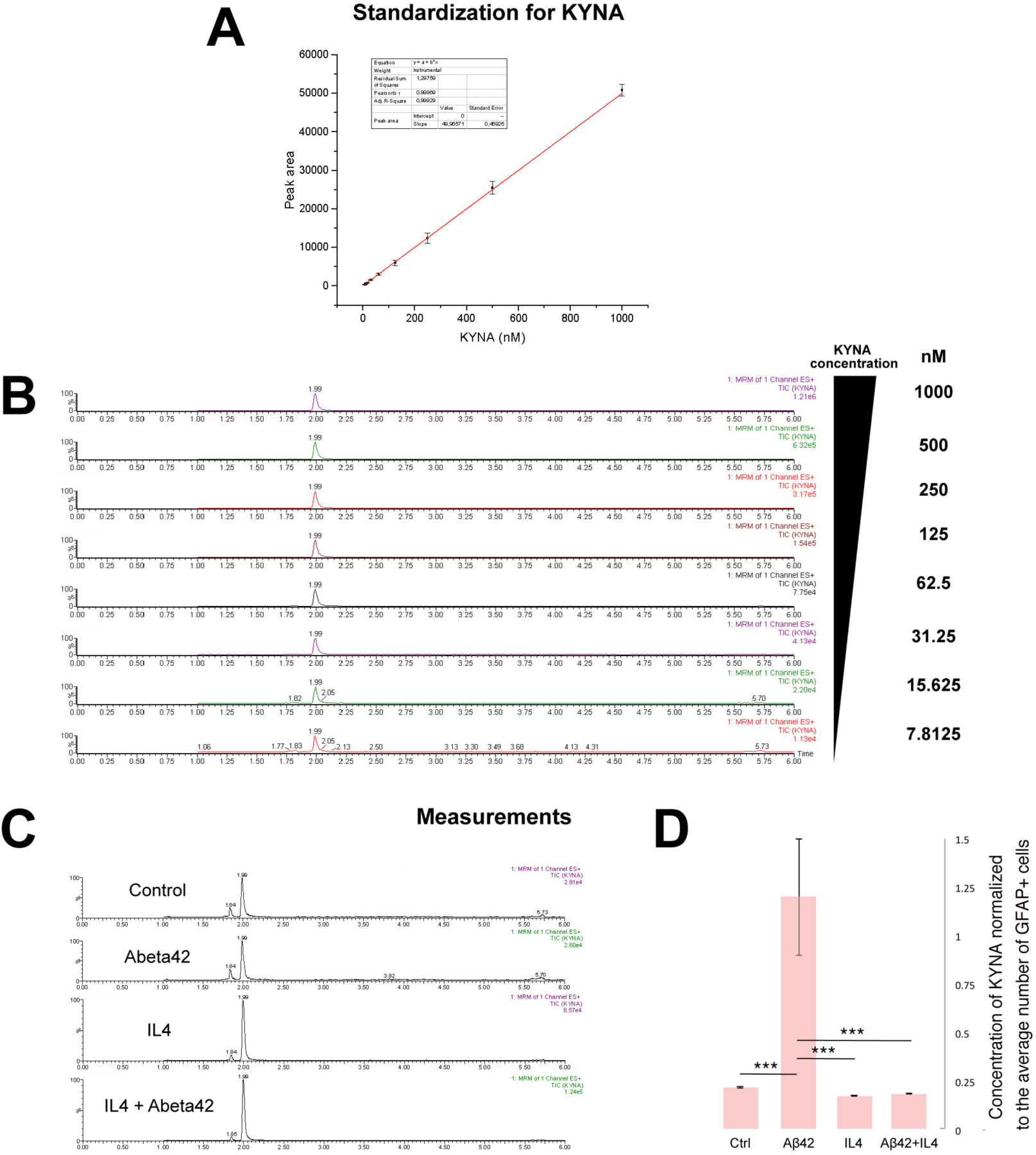
Measurement of KYNA concentration. (A) Calibration curve for various concentrations of KYNA. (B) Liquid chromatography-mass spectroscopy (LC-MS/MS) running curves for different KYNA concentrations. (C) LC-MS/MS running curves for untreated (control), Aβ42-treated, IL4-treated, and Aβ42+IL4-treated gels. (D) Normalized quantification for the concentrations of KYNA in untreated (control), Aβ42-treated, IL4-treated, and Aβ42+IL4-treated gels. All gels are 3 weeks of culture. Related to Figure 3.

**Supplementary Figure 4.**
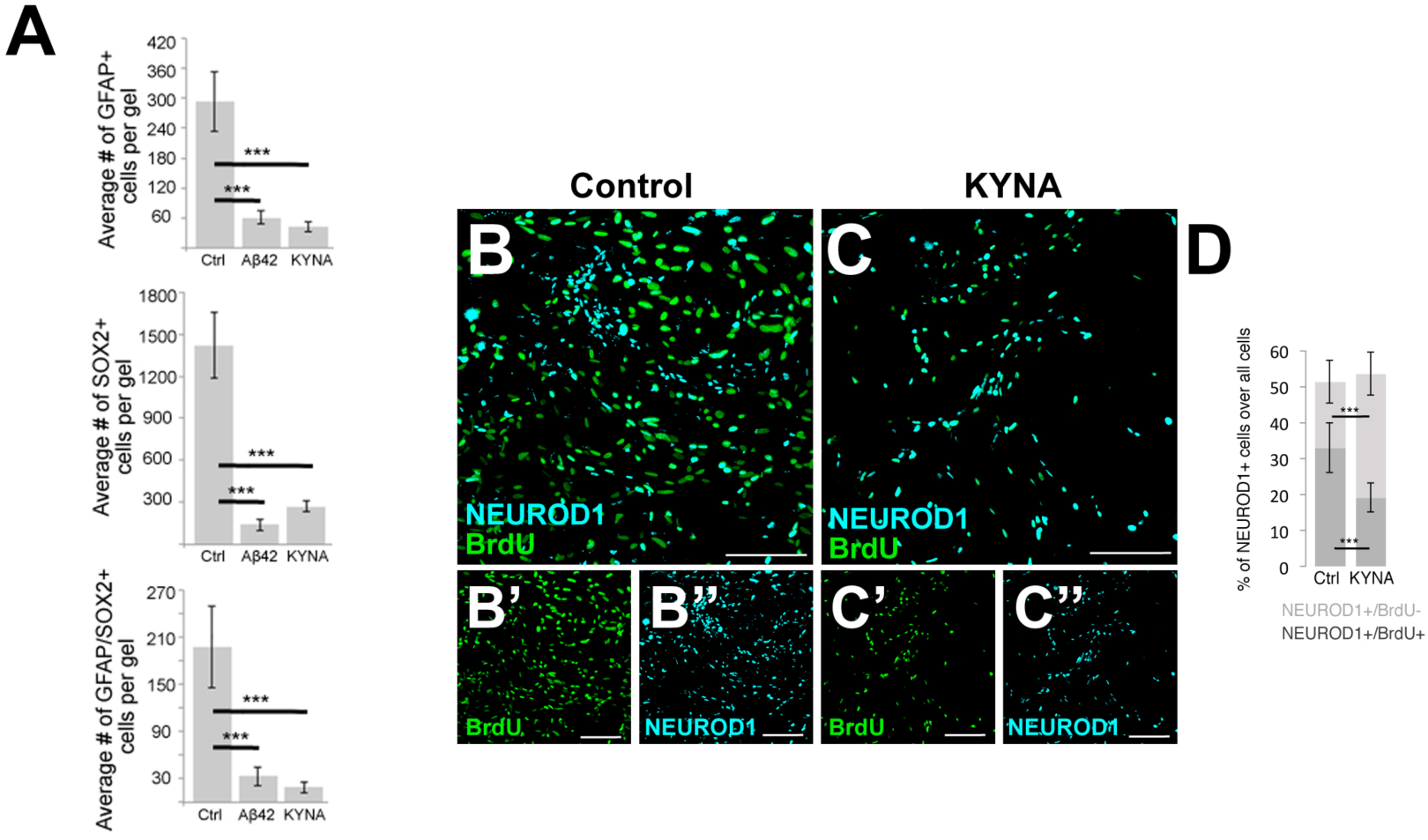
Effects of Aβ42 and KYNA on the number of NSCs, and the effects of Kynurenic acid on neurogenic capacity. (A) Quantification graphs for the number of GFAP+/SOX2-, GFAP-/SOX2+ and GFAP+/SOX2+ progenitors in control, Aβ42-treated and KYNA-treated gels. (B, C’’) NEUROD1 and BrdU in control (B) and Kynurenic acid (KYNA)-treated gels (C). Smaller images under the panels are single fluorescent images (B’,B’’, C’, C’’). (D) Quantification graph for B-C’’. Scale bars: 100 *μ*m. All gels are 3 weeks of culture. Related to Figure 3 and 4.

